# Identification of a High-frequency Intra-host SARS-CoV-2 spike Variant with Enhanced Cytopathic and Fusogenic Effect

**DOI:** 10.1101/2020.12.03.409714

**Authors:** Lynda Rocheleau, Geneviève Laroche, Kathy Fu, Corina M Stewart, Abdulhamid O Mohamud, Marceline Côté, Patrick M Giguère, Marc-André Langlois, Martin Pelchat

**Affiliations:** Department of Biochemistry, Microbiology and Immunology, Faculty of Medicine, University of Ottawa, Ottawa, Ontario, K1H 8M5, Canada; University of Ottawa Brain and Mind Research Institute, University of Ottawa, 451 Smyth Road, Ottawa, Ontario K1H 8M5, Canada; uOttawa Center for Infection, Immunity and Inflammation (CI3), Ottawa, Canada; Ottawa Institute of Systems Biology, University of Ottawa, Ottawa, Canada

**Keywords:** COVID-19, SARS-CoV-2, syncytia, genetic variants, high-throughput sequencing, Spike protein

## Abstract

The severe acute respiratory syndrome coronavirus 2 (SARS-CoV-2) is a virus that is continuously evolving. Although its RNA-dependent RNA polymerase exhibits some exonuclease proofreading activity, viral sequence diversity can be produced by replication errors and host factors. A diversity of genetic variants can be observed in the intra-host viral population structure of infected individuals. Most mutations will follow a neutral molecular evolution and won’t make significant contributions to variations within and between infected hosts. Herein, we profiled the intra-sample genetic diversity of SARS-CoV-2 variants using high-throughput sequencing datasets from 15,289 infected individuals and infected cell lines. Most of the genetic variations observed, including C->U and G->U, were consistent with errors due to heat-induced DNA damage during sample processing and/or sequencing protocols. Despite high mutational background, we identified recurrent intra-variable positions in the samples analyzed, including several positions at the end of the gene encoding the viral Spike (S) protein. Strikingly, we observed a high-frequency C->A missense mutations resulting in the S protein lacking the last 20 amino acids (SΔ20). We found that this truncated S protein undergoes increased processing and increased syncytia formation, presumably due to escaping M protein retention in intracellular compartments. Our findings suggest the emergence of a high-frequency viral sublineage that is not horizontally transmitted but potentially involved in intra-host disease cytopathic effects.

**IMPORTANCE:** The mutation rate and evolution of RNA viruses correlate with viral adaptation. While most mutations do not have significant contributions to viral molecular evolution, some are naturally selected and cause a genetic drift through positive selection. Many recent SARS-CoV-2 variants have been recently described and show phenotypic selection towards more infectious viruses. Our study describes another type of variant that does not contribute to inter-host heterogeneity but rather phenotypic selection toward variants that might have increased cytopathic effects. We identified that a C-terminal truncation of the Spike protein removes an important ER-retention signal, which consequently results in a Spike variant that easily travels through the Golgi toward the plasma membrane in a pre-activated conformation, leading to increased syncytia formation.

## INTRODUCTION

Observed for the first time in 2019, the severe acute respiratory syndrome coronavirus 2 (SARS-CoV-2) and its associated disease, COVID-19, has caused significant worldwide mortality and unprecedented economic burdens. SARS-CoV-2 is an enveloped virus with a non-segmented, positive-sense single-stranded RNA (vRNA) genome comprised of a ∼30K nucleotides (1, 2). The virus is composed of four main structural proteins, encoded in the last 3’ terminal third of the viral genome: the spike glycoprotein (S), membrane (M), envelope (E) and the nucleocapsid (N) (3–5). Attachment to the host receptor angiotensin-converting enzyme 2 (ACE2) is mediated by the S protein expressed on the surface of the virion (6). Following its association, the S protein is cleaved into two separate polypeptides (S1 and S2), which triggers the fusion of the viral particle with the cellular membrane (7, 8). Once inside a cell, its RNA-dependent RNA polymerase (RdRp), which is encoded in the first open reading frame of the viral genome (9), carries out transcription and replication of the vRNA genome. In addition, mRNAs coding for the structural proteins (*e*.*g*., S, M, E and N) are expressed by subgenomic RNAs (9). Once translated, the S, M and E proteins localize and accumulate at the CoV budding site in the endoplasmic reticulum–Golgi intermediate compartment (ERGIC) (10). One aspect of CoV biology is that CoV virions bud into the lumen of the secretory pathway at the ERGIC and must then traffic through the Golgi complex and anterograde system to be efficiently released from host cells (11). The S protein possesses an endoplasmic reticulum retrieval signal (ERRS) at its carboxy terminus, which is required for trafficking through the ERGIC (12). At this location, the spike protein interacts with the M protein, which has been shown to be essential for accumulation at the ERGIC. The N protein then associates with the viral genome and assembles into virions, which are transported along the endosomal network and released by exocytosis (9). If not retained at ERGIC, the S proteins traffics through the Golgi and is pre-activated by resident proteases prior to reaching the plasma membrane. Here it can mediate cell fusion between adjacent cells, resulting in the production of multinucleated cells or syncytia (8, 13, 14). Genomic sequencing of SARS-CoV-2 vRNA from infected populations has demonstrated genetic heterogeneity (15–21). Several recurrent mutations have been identified in consensus sequences, and the geographical distribution of clades was established. Because they induce an abundance of missense rather than synonymous or non-sense mutations, it was suggested that regions of the SARS-CoV-2 genome were actively evolving and might contribute to pandemic spreading (21). It was observed that variations are mainly comprised of transition mutations (purine->purine or pyrimidine->pyrimidine) with a prevalence of C->U transitions and might occur within a sequence context reminiscent of APOBEC-mediated deamination (*i*.*e*., [AU]C[AU]; (22, 23)). Consequently, it was proposed that host editing enzymes might be involved in coronavirus genome editing (24, 25).

Consensus mutations are only part of the genetic landscape with regards to RNA viruses. Replication of RNA viruses typically produces quasispecies in which the viral RNA genomes do not exist as single sequence entity but rather as a population of genetic variants (26). These mutations are most frequently caused by both the error-prone nature of each of their respective viral RdRps, as well as by host RNA editing enzymes such as APOBECs and ADARs (27). However, the RdRp complex of large RNA viruses, such as coronaviruses, sometimes possess exonuclease proofreading activity, and consequently have lower error rates (26, 28). Quasispecies may sometimes exhibit diminished replicative fitness or deleterious mutations and exert different roles that are not directly linked to viral genomic propagation (29). Mutations that form the intra-host genetic spectrum have been shown to help viruses evade cytotoxic T cell recognition and neutralizing antibodies, rendering these viruses more resistant to antiviral drugs (29). Additionally, these mutations can also be involved in modulating the virulence and transmissibility of the quasispecies (29).

In this study, we focussed on assessing intra-host genetic variations of SARS-CoV-2. We analyzed high-throughput sequencing datasets to profile the sequence diversity of SARS-CoV-2 variants within distinct sample populations. We observed high genetic intra-variability of the viral genome. By comparing variation profiles between samples from different donors and cell lines, we identified highly conserved subspecies that independently and recurrently arose in different datasets and, therefore, in different individuals. We further analyzed the dominant variant SΔ20 in a functional assay and demonstrate that this truncated S protein avoids inhibition caused by M protein and enhances syncytium formation. We provide evidence for the existence of a consistently emerging variant identified across geographical regions that may influence intra-host SARS-CoV-2 pathogenicity.

## MATERIAL AND METHODS

### Analysis of intra-variability within SARS-CoV-2 samples

15,289 publicly available high-throughput sequencing datasets were downloaded from the NCBI Sequence Read Archive (up to July 10, 2020). They comprise of 15,224 and 65 datasets from infected individuals and infected cell lines, respectively. The datasets from infected cells were generated by Blanco-Melo et al. (30). Duplicated reads were combined to reduce amplification bias and mapped to the SARS-CoV-2 isolate Wuhan-Hu-1 reference genome (NC_045512v2) using hisat2 (v.2.1.0)(31). For each dataset, the consensus sequences and the frequency of nucleotides at each position were extracted from files generated by bcftools (v.1.10.2) of the samtools package (v.1.1) with an in-house Perl script (32, 33). All further calculations were performed in R. To reduce the number of variations due to sequencing errors and/or protocol differences, only positions mapped with a sequencing depth of 50 reads and having at least 5 reads with variations compared to the sample consensus were considered. Sequence logos were generated with the ggseqlogo package (v.0.1) (34).

### Differential expression analysis of transcript coding for APOBECs and ADARs

High-throughput sequencing datasets generated in a recent transcription profiling study of several cell lines infected with SARS-CoV-2 were downloaded from SRA (30). Duplicated reads were combined and mapped to the human reference genome (Homo_sapiens.GRCh38.83) using hisat2 (v.2.1.0)(31). Transcript abundance was performed using HTSeq 0.12.4 (35) and normalized into Transcripts Per Million (TPM) in R.

### Cell culture and plasmids

Human embryonic kidney 293T (HEK293T) were obtained from the American Type Culture Collection (ATCC CRL-11268) and maintained in Dulbecco’s Modified Eagle’s Medium (DMEM) supplemented with 5% fetal bovine serum (Fisher Scientific), 5% bovine calf serum (Fisher Scientific) and 100 U/mL penicillin, 100 µg/mL streptomycin (Fisher Scientific). HEK293T stably expressing human ACE2 (HEK-293T-hACE2 cell line, bei RESOURCES) were cultured and maintained in DMEM (Corning) supplemented with 10% fetal bovine serum (Sigma), 100 U/mL penicillin and 100 µg/mL streptomycin. All cells were cultured at 37 °C in a humidified atmosphere containing 5% CO_2_. pCAGGS expressing the SARS-CoV-2 S protein (Wuhan-Hu-1; WT) was provided by Dr. Florian Krammer (Mount Sinai). SARS-CoV-2 SΔ20 (SΔ20) was generated using overlapping PCR to introduce a termination codon at residue 1254. The expression construct encoding SARS-CoV-2 M was generated by PCR amplification of the M gene from pLVX-EF1alpha-SARS-CoV-2-M-2xStrep-IRES-Puro (kind gift of Dr. Nevan Krogan, UCSF) and addition of a stop codon to remove the Strep tag prior to cloning into pCAGGS.

### Syncytium Formation Assay

HEK-293T-hACE2 cells were seeded in 24-well plates in complete media to obtain a 90% confluence the following day. Cells were then transiently co-transfected using JetPRIME (Polyplus Transfection, France) with plasmids encoding GFP (MLV-GFP, a kind gift of Dr. James Cunningham, Brigham and Women’s Hospital), SARS-CoV-2 S or SARS-CoV-2 SΔ20, and M or pCAGGS at a 0.15:0.2:0.65 ratio. 18 hours post-transfection, cells were imaged (ZOE Fluorescent Cell Imager, Bio-Rad) for syncytia formation using the Green channel to visualize fusion of GFP positive cells as performed previously (36).

### Western Blot analysis

HEK293T cells were transfected with the empty vector (pCAGGS), or with SARS-CoV-2 S or SARS-CoV-2 SΔ20 and M or pCAGGS using JetPRIME at a 1:1 ratio. The following day, cells were washed once with cold PBS and lysed in cold lysis buffer (1% Triton X-100, 0.1% IGEPAL CA-630, 150mM NaCl, 50mM Tris-HCl, pH 7.5) containing protease and phosphatase inhibitors (Cell Signaling). Proteins in cell lysates were resolved on 4-12% gradient SDS-polyacrylamide gels (NuPage, Invitrogen) and transferred to polyvinylidenedifluoride (PVDF) membranes. Membranes were blocked for 1h at RT with blocking buffer (5% skim milk powder dissolved in 25mM Tris, pH 7.5, 150mM NaCl, and 0.1% Tween-20 [TBST]). Processing of spike was detected by immunoblotting using an anti-S1 (SARS-CoV/SARS-CoV-2 spike protein S1 polyclonal, Invitrogen) and anti-S2 antibody (SARS-CoV/SARS-CoV-2 spike protein S2 monoclonal, Invitrogen). Overexpression of M was also detected by immunoblotting and using an anti-M antibody (Rabbit anti-SARS Membrane protein, NOVUS). Membranes were incubated overnight at 4°C with the appropriate primary antibody in the blocking buffer. Blots were then washed in TBST and incubated with HRP-conjugated secondary antibody for 1h at room temperature (anti-mouse HRP (Cell Signaling) and anti-rabbit HRP (Cell Signaling). Membranes were washed, incubated in chemiluminescence substrate (SuperSignal West Femto Maximum Sensitivity Substrate, ThermoFisher scientific), and imaged using the ChemiDoc XRS+ imaging system (Bio-Rad). In some instances, the same membrane was stripped and re-probed for actin (Monoclonal Anti-β-Actin, Millipore Sigma).

## RESULTS

### High intra-genetic variability of the SARS-CoV-2 genome in infected individuals

To assess the extent of SARS-CoV-2 sequence intra-genetic variability, we analyzed 15,224 publicly available high-throughput sequencing datasets from infected individuals. The raw sequencing reads were mapped to the SARS-CoV-2 isolate Wuhan-Hu-1 reference genome, and the composition of each nucleotide at each position on the viral genome was generated. Consensus sequences were produced for each dataset and the nucleotide composition for each position were compared to respective consensuses. To reduce the number of variations due to amplification bias and sequencing errors, duplicated reads were combined, and only positions mapped with a sequencing depth of 50 reads and having at least 5 reads with variations compared to the sample consensus were considered. Overall, we identified 301,742 variations from 11,362 samples located on 26,113 positions of the 29,903 nt SARS-CoV-2 genome. We observed an average of 26.6+/-132.0 variable nucleotides per sample (ranging from 1 to 5295 variations/sample; Fig. 1A).

**Fig. 1:**
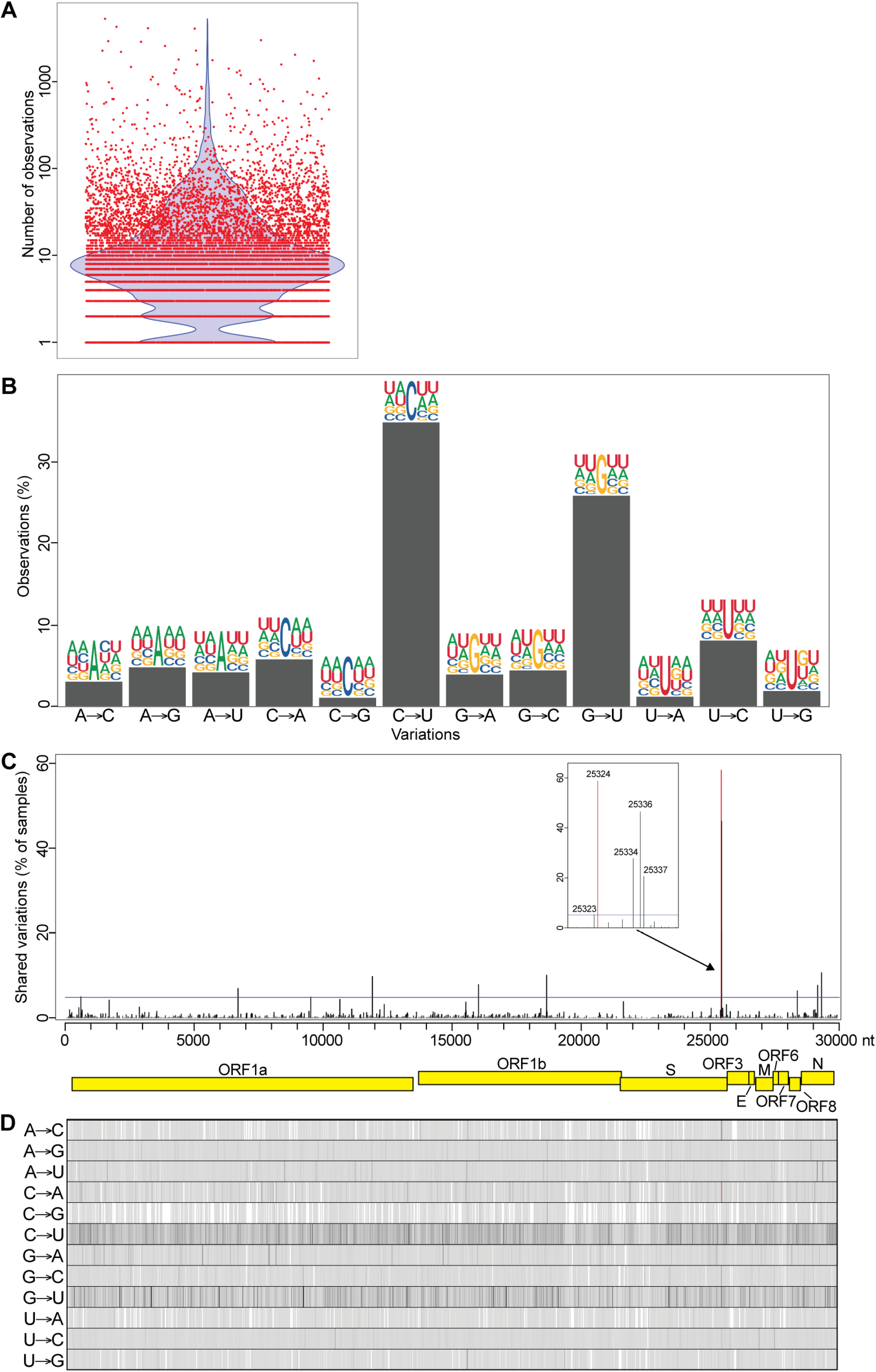
Intra-sample variability of the SARS-CoV-2 genome in infected individuals. (**A**) Number of intra-variations observed for each sample analyzed. The red dots represent the 11,362 samples analyzed and the blue violon shows the distribution of the data. (**B**) Type of variation and sequence context for each intra-sample variable position. Bars represent the percentage of each type. Sequence context is represented by logos comprised of the consensus nucleotides (center) with two nucleotides upstream and two downstream from each intra-sample variable position. (**C**) Recurrent intra-genetic variations are represented as percentages of samples containing variations at each position. The SARS-CoV-2 genome and its genes are represented by yellow boxes below the graph. The blue line indicates 5% shared variations and was used to extract the recurrent intra-sample variations listed in Table 1. The inset represents a magnification of the cluster identified at the end of the S gene. (**D**) One-dimension representation of the data shown in panel C for each type of variation individually. The location of the C->A variation at position 25,324 is indicated by a red line in (C) and (D).

### Analysis of the type of intra-genetic variations present in SARS-CoV-2 samples from infected individuals

The analysis of the type of nucleotide changes within samples revealed that 52.2% were transitions (either purine->purine or pyrimidine->pyrimidine) and 47.8% were transversions (purine->pyrimidine or pyrimidine->purine). Notably, the highest nucleotide variations corresponded to C->U transitions (43.5%) followed by G->U transversion (28.1%; Fig. 1B), both types encompassing 71.6% of all variations. Since editing by host enzymes depends on the sequence context, we extracted two nucleotides upstream and downstream from each genomic position corresponding to variations and generated sequence logos. Our results indicated a high number of As and Us around all variation types and sites (62.1+/-3.4%; Fig. 1B). Because SARS-CoV-2 is composed of 62% A/U, this suggests that the observed number of As and Us around variation sites are mainly due to the A/U content of the viral genome, the fact that no motifs are enriched around these sites, and that these intra-genetic variations are likely not originating from host editing enzymes.

### Identification of recurrent genetic variants of SARS-CoV-2 in samples from infected individuals

To identify biologically relevant intra-genetic variations, we examined the variable positions that are recurrent in the samples analyzed. The variable positions were tabulated for each sample and then recurrent intra-genetic variations were calculated as percentages of samples containing variation at each position. Most variations are distributed homogeneously on the viral genome and most are poorly shared amongst samples (Fig. 1C and 1D). However, our analysis reveals 15 recurrent intra-variations shared by at least 5% of the samples analyzed (Fig. 1C, above blue line; Table 1). Amongst these, four transversions (at nt 25,324, 25,334, 25,336 and 25,337) located at the 3’ end of the S gene are the most recurrent variations (inset of Fig. 1C and Table 1). Three of these transversions (at nt 25,334, 25,336 and 25,337) correspond to missense mutations: E1258D (46.4%), E1258Q (27.6%) and D1259H (20.1%). Interestingly, the most observed variation (at nt 25,324) is shared by 58.7% of the samples (6,668 of the 11,362 samples) and corresponds to a C->A transversion producing a nonsense mutation at amino acid 1,254 of the S protein (Fig. 1C and 1D, red line; Fig. 2B, red rectangle). The resulting S protein lacks the last 20 amino acids (SΔ20), which includes the ERRS motif at its carboxy terminus (Fig. 2B, white letters on a black background). Amongst the sample with this intra-genetic variation, this C->A transversion represents from 2.9% to 42.4% of the subspecies identified (mean or 8.2+/-2.9%; Fig. 2C and Table 1).

**Table 1:**
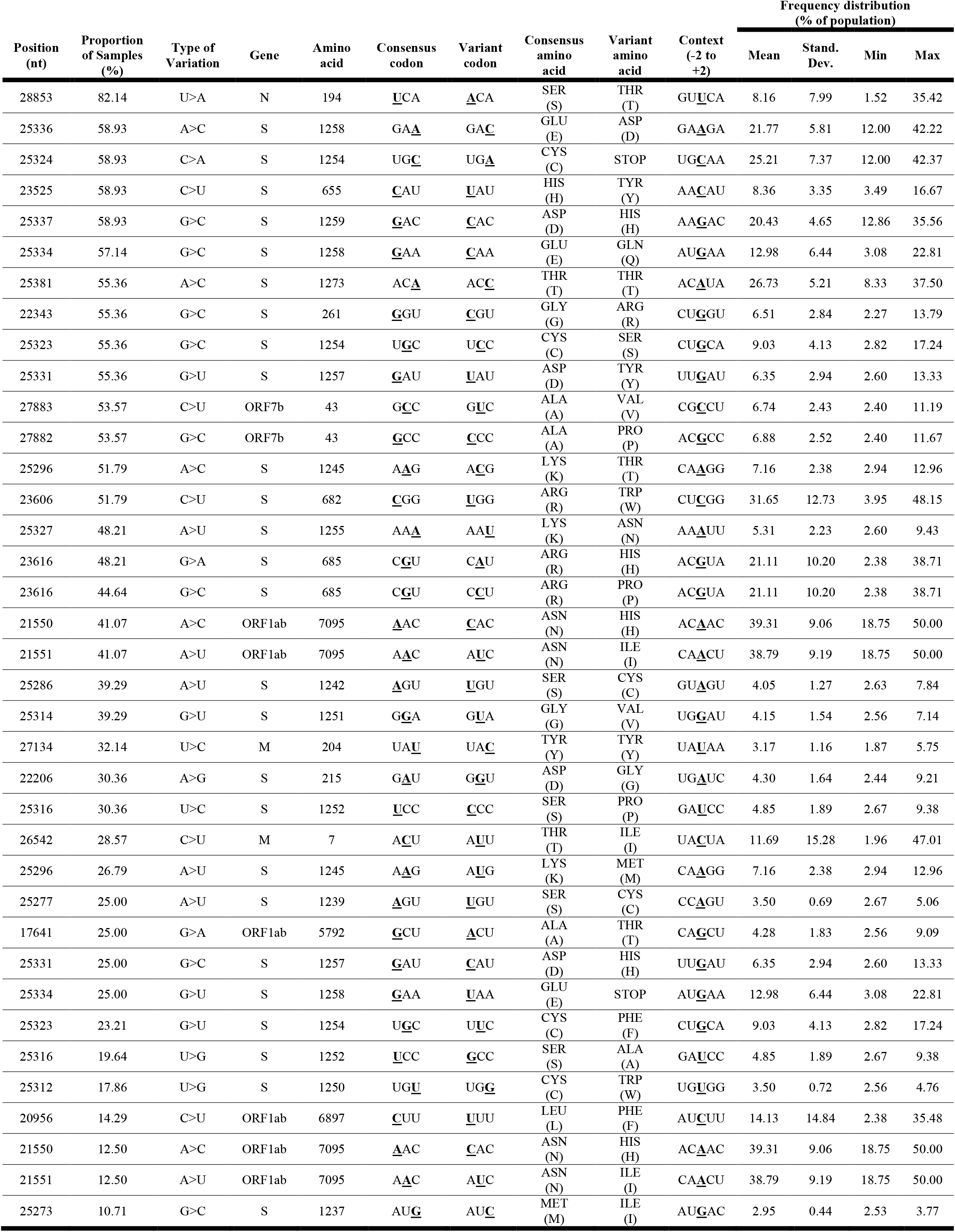
Recurrent SARS-CoV-2 genome intra-variations shared by at least 5% infected individuals. Frequency distributions were calculated using data similar to Fig. 2C. The variations are sorted by their recurrence, with the most shared variation at the top of the table.

| Position<br>(nt) | Proportion<br>of Samples<br>(%) | Type of<br>Variation | Gene | Amino<br>acid | Consensus<br>codon | Variant<br>codon | Consensus<br>amino<br>acid | Variant<br>amino<br>acid | Context<br>(-2 to<br>+2) | Frequency distribution<br>(% of population) |  |  |  |
| --- | --- | --- | --- | --- | --- | --- | --- | --- | --- | --- | --- | --- | --- |
|  |  |  |  |  |  |  |  |  |  | Mean | Stand.<br>Dev. | Min | Max |
| 28853 | 82.14 | U>A | N | 194 | <u>U</u>CA | <u>A</u>CA | SER<br>(S) | THR<br>(T) | GU<u>U</u>CA | 8.16 | 7.99 | 1.52 | 35.42 |
| 25336 | 58.93 | A>C | S | 1258 | GA<u>A</u> | GAC | GLU<br>(E) | ASP<br>(D) | GA<u>A</u>GA | 21.77 | 5.81 | 12.00 | 42.22 |
| 25324 | 58.93 | C>A | S | 1254 | UG<u>C</u> | UG<u>A</u> | CYS<br>(C) | STOP | UGCAA | 25.21 | 7.37 | 12.00 | 42.37 |
| 23525 | 58.93 | C>U | S | 655 | <u>C</u>AU | <u>U</u>AU | HIS<br>(H) | TYR<br>(Y) | AA<u>C</u>AU | 8.36 | 3.35 | 3.49 | 16.67 |
| 25337 | 58.93 | G>C | S | 1259 | <u>G</u>AC | <u>C</u>AC | ASP<br>(D) | HIS<br>(H) | AA<u>G</u>AC | 20.43 | 4.65 | 12.86 | 35.56 |
| 25334 | 57.14 | G>C | S | 1258 | <u>G</u>AA | <u>C</u>AA | GLU<br>(E) | GLN<br>(Q) | AU<u>G</u>AA | 12.98 | 6.44 | 3.08 | 22.81 |
| 25381 | 55.36 | A>C | S | 1273 | AC<u>A</u> | ACC | THR<br>(T) | THR<br>(T) | AC<u>A</u>UA | 26.73 | 5.21 | 8.33 | 37.50 |
| 22343 | 55.36 | G>C | S | 261 | <u>G</u>GU | <u>C</u>GU | GLY<br>(G) | ARG<br>(R) | CU<u>G</u>GU | 6.51 | 2.84 | 2.27 | 13.79 |
| 25323 | 55.36 | G>C | S | 1254 | UG<u>C</u> | UCC | CYS<br>(C) | SER<br>(S) | CUGCA | 9.03 | 4.13 | 2.82 | 17.24 |
| 25331 | 55.36 | G>U | S | 1257 | <u>G</u>AU | <u>U</u>AU | ASP<br>(D) | TYR<br>(Y) | UU<u>G</u>AU | 6.35 | 2.94 | 2.60 | 13.33 |
| 27883 | 53.57 | C>U | ORF7b | 43 | GCC | GU<u>C</u> | ALA<br>(A) | VAL<br>(V) | CGCCU | 6.74 | 2.43 | 2.40 | 11.19 |
| 27882 | 53.57 | G>C | ORF7b | 43 | <u>G</u>CC | <u>C</u>CC | ALA<br>(A) | PRO<br>(P) | AC<u>G</u>CC | 6.88 | 2.52 | 2.40 | 11.67 |
| 25296 | 51.79 | A>C | S | 1245 | A<u>A</u>G | ACG | LYS<br>(K) | THR<br>(T) | CA<u>A</u>GG | 7.16 | 2.38 | 2.94 | 12.96 |
| 23606 | 51.79 | C>U | S | 682 | <u>C</u>GG | <u>U</u>GG | ARG<br>(R) | TRP<br>(W) | CUCGG | 31.65 | 12.73 | 3.95 | 48.15 |
| 25327 | 48.21 | A>U | S | 1255 | AA<u>A</u> | AA<u>U</u> | LYS<br>(K) | ASN<br>(N) | AA<u>A</u>UU | 5.31 | 2.23 | 2.60 | 9.43 |
| 23616 | 48.21 | G>A | S | 685 | <u>C</u>GU | <u>C</u>AU | ARG<br>(R) | HIS<br>(H) | AC<u>G</u>UA | 21.11 | 10.20 | 2.38 | 38.71 |
| 23616 | 44.64 | G>C | S | 685 | <u>C</u>GU | <u>C</u>CU | ARG<br>(R) | PRO<br>(P) | AC<u>G</u>UA | 21.11 | 10.20 | 2.38 | 38.71 |
| 21550 | 41.07 | A>C | ORF1ab | 7095 | <u>A</u>AC | <u>C</u>AC | ASN<br>(N) | HIS<br>(H) | AC<u>A</u>AC | 39.31 | 9.06 | 18.75 | 50.00 |
| 21551 | 41.07 | A>U | ORF1ab | 7095 | A<u>A</u>C | A<u>U</u>C | ASN<br>(N) | ILE<br>(I) | CA<u>A</u>CU | 38.79 | 9.19 | 18.75 | 50.00 |
| 25286 | 39.29 | A>U | S | 1242 | <u>A</u>GU | <u>U</u>GU | SER<br>(S) | CYS<br>(C) | GU<u>A</u>GU | 4.05 | 1.27 | 2.63 | 7.84 |
| 25314 | 39.29 | G>U | S | 1251 | <u>G</u>GA | <u>G</u>UA | GLY<br>(G) | VAL<br>(V) | UG<u>G</u>AU | 4.15 | 1.54 | 2.56 | 7.14 |
| 27134 | 32.14 | U>C | M | 204 | UA<u>U</u> | UAC | TYR<br>(Y) | TYR<br>(Y) | UA<u>U</u>AA | 3.17 | 1.16 | 1.87 | 5.75 |
| 22206 | 30.36 | A>G | S | 215 | <u>G</u>AU | <u>G</u>GU | ASP<br>(D) | GLY<br>(G) | UG<u>A</u>UC | 4.30 | 1.64 | 2.44 | 9.21 |
| 25316 | 30.36 | U>C | S | 1252 | <u>U</u>CC | <u>C</u>CC | SER<br>(S) | PRO<br>(P) | GA<u>U</u>CC | 4.85 | 1.89 | 2.67 | 9.38 |
| 26542 | 28.57 | C>U | M | 7 | A<u>C</u>U | A<u>U</u>U | THR<br>(T) | ILE<br>(I) | UA<u>C</u>UA | 11.69 | 15.28 | 1.96 | 47.01 |
| 25296 | 26.79 | A>U | S | 1245 | A<u>A</u>G | A<u>U</u>G | LYS<br>(K) | MET<br>(M) | CA<u>A</u>GG | 7.16 | 2.38 | 2.94 | 12.96 |
| 25277 | 25.00 | A>U | S | 1239 | <u>A</u>GU | <u>U</u>GU | SER<br>(S) | CYS<br>(C) | CC<u>A</u>GU | 3.50 | 0.69 | 2.67 | 5.06 |
| 17641 | 25.00 | G>A | ORF1ab | 5792 | <u>G</u>CU | <u>A</u>CU | ALA<br>(A) | THR<br>(T) | CAGCU | 4.28 | 1.83 | 2.56 | 9.09 |
| 25331 | 25.00 | G>C | S | 1257 | <u>G</u>AU | <u>C</u>AU | ASP<br>(D) | HIS<br>(H) | UU<u>G</u>AU | 6.35 | 2.94 | 2.60 | 13.33 |
| 25334 | 25.00 | G>U | S | 1258 | <u>G</u>AA | <u>U</u>AA | GLU<br>(E) | STOP | AU<u>G</u>AA | 12.98 | 6.44 | 3.08 | 22.81 |
| 25323 | 23.21 | G>U | S | 1254 | UG<u>C</u> | U<u>U</u>C | CYS<br>(C) | PHE<br>(F) | CUGCA | 9.03 | 4.13 | 2.82 | 17.24 |
| 25316 | 19.64 | U>G | S | 1252 | <u>U</u>CC | <u>G</u>CC | SER<br>(S) | ALA<br>(A) | GA<u>U</u>CC | 4.85 | 1.89 | 2.67 | 9.38 |
| 25312 | 17.86 | U>G | S | 1250 | UG<u>U</u> | UG<u>G</u> | CYS<br>(C) | TRP<br>(W) | UG<u>U</u>GG | 3.50 | 0.72 | 2.56 | 4.76 |
| 20956 | 14.29 | C>U | ORF1ab | 6897 | <u>C</u>UU | <u>U</u>UU | LEU<br>(L) | PHE<br>(F) | AU<u>C</u>UU | 14.13 | 14.84 | 2.38 | 35.48 |
| 21550 | 12.50 | A>C | ORF1ab | 7095 | <u>A</u>AC | <u>C</u>AC | ASN<br>(N) | HIS<br>(H) | AC<u>A</u>AC | 39.31 | 9.06 | 18.75 | 50.00 |
| 21551 | 12.50 | A>U | ORF1ab | 7095 | A<u>A</u>C | A<u>U</u>C | ASN<br>(N) | ILE<br>(I) | CA<u>A</u>CU | 38.79 | 9.19 | 18.75 | 50.00 |
| 25273 | 10.71 | G>C | S | 1237 | AU<u>G</u> | AU<u>C</u> | MET<br>(M) | ILE<br>(I) | AU<u>G</u>AC | 2.95 | 0.44 | 2.53 | 3.77 |

**Fig. 2:**
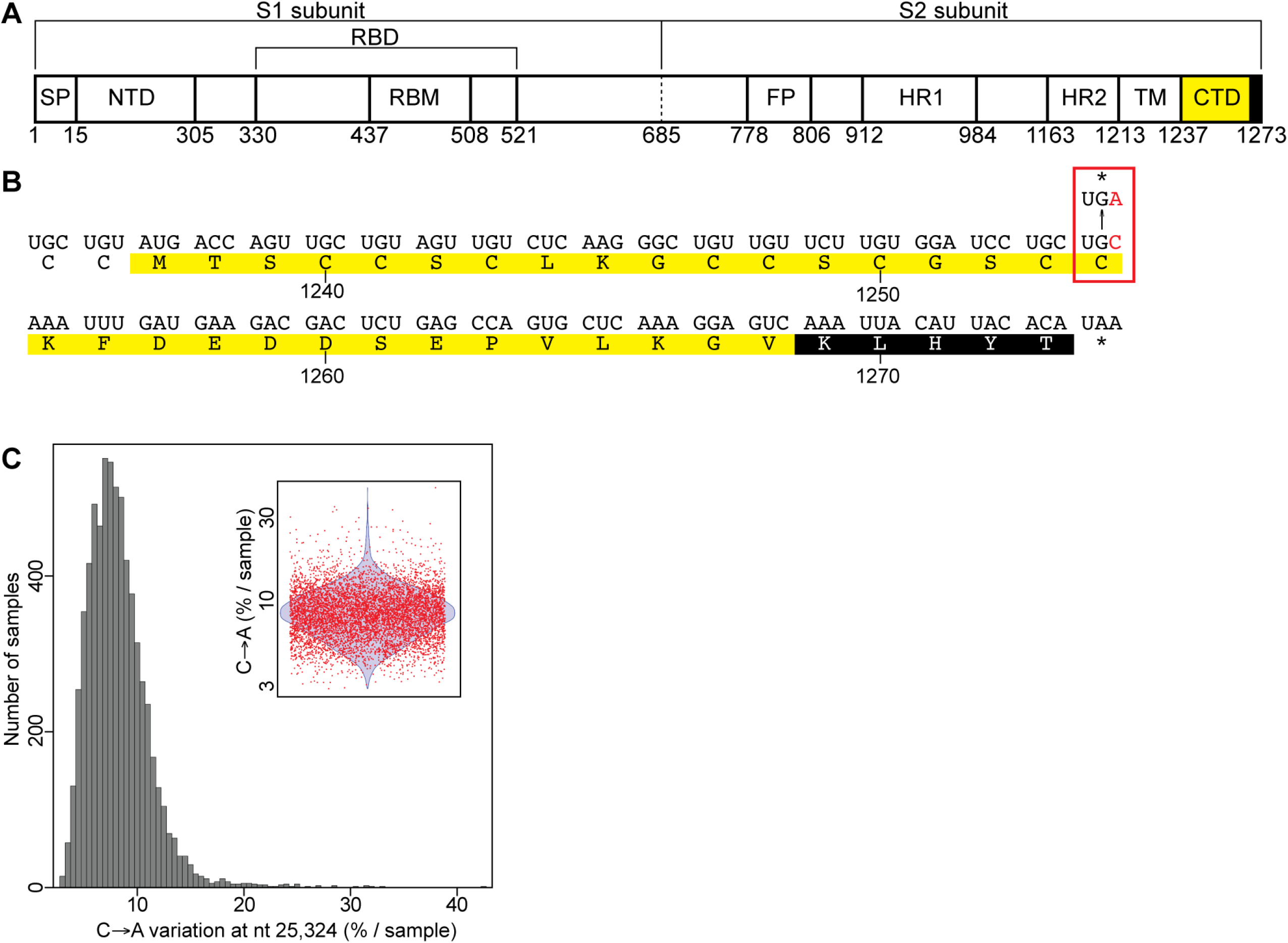
Localization of the C->A missense mutation on the SARS-CoV-2 S protein. (**A**) Schematic representation of the functional domain of the SARS-CoV-2 S protein. (**B**) Localization of the C->A variation on the carboxy terminal domain (CTD) of the S protein. The mutation is colored and boxed in red. The carboxy terminal domain (CTD) and the ERRS are colored in yellow and black, respectfully in (A) and (B). (**C**) Distribution of the intra-sample proportion of the C->A transversion at position 25,324 in the 6,668 samples containing this subspecies. The inset represents the distribution using red dots to represent the samples having this intra-genetic variation and a blue violon to show the distribution of the data.

### Analysis of intra-genetic variations present in SARS-CoV-2 samples from infected cells

To further investigate variations in a more controlled system and to determine whether host proteins are involved in SARS-CoV-2 genome editing, we used 65 high-throughput sequencing datasets generated in a recent transcription profiling study of several cell lines infected with SARS-CoV-2 (30). Firstly, we mapped raw sequencing reads to the human genome to assess host modifying enzyme expression. For all cell lines, normalized counts for mRNAs corresponding to most host modifying enzymes were very low or non-detected (Fig. 3), suggesting that these cell lines poorly expressed these host editing proteins. As above, the raw sequencing reads from infected cells were mapped to the SARS-CoV-2 genome sequence, the composition of each nucleotide at each position on the viral genome were generated, and nucleotide variations compared to respective consensus sequences were calculated. Because the sequencing depths of the samples were low, we considered positions mapped by at least 20 reads and having at least 2 reads with variation compared to the sample consensus. In the samples derived from infected cells, we observed 29.7% and 70.3% of transitions and transversions, respectively. Similar to observations in samples from infected individuals, the highest nucleotide variations corresponded to G->U transversions (26.1%) and C->U transitions (21.6%; Fig. 4B). We then analyzed nucleotide compositions two nucleotides upstream and downstream of the intra-genetic variations. As above, a high number of A/U (57.8+/-7.7%) were present around variation sites (Fig. 4B), consistent with the 62% A/U composition of the SARS-CoV-2 genome, indicating no enrichment of sequence motifs around these sites, except for the expected high number of As and Us.

**Fig. 3:**
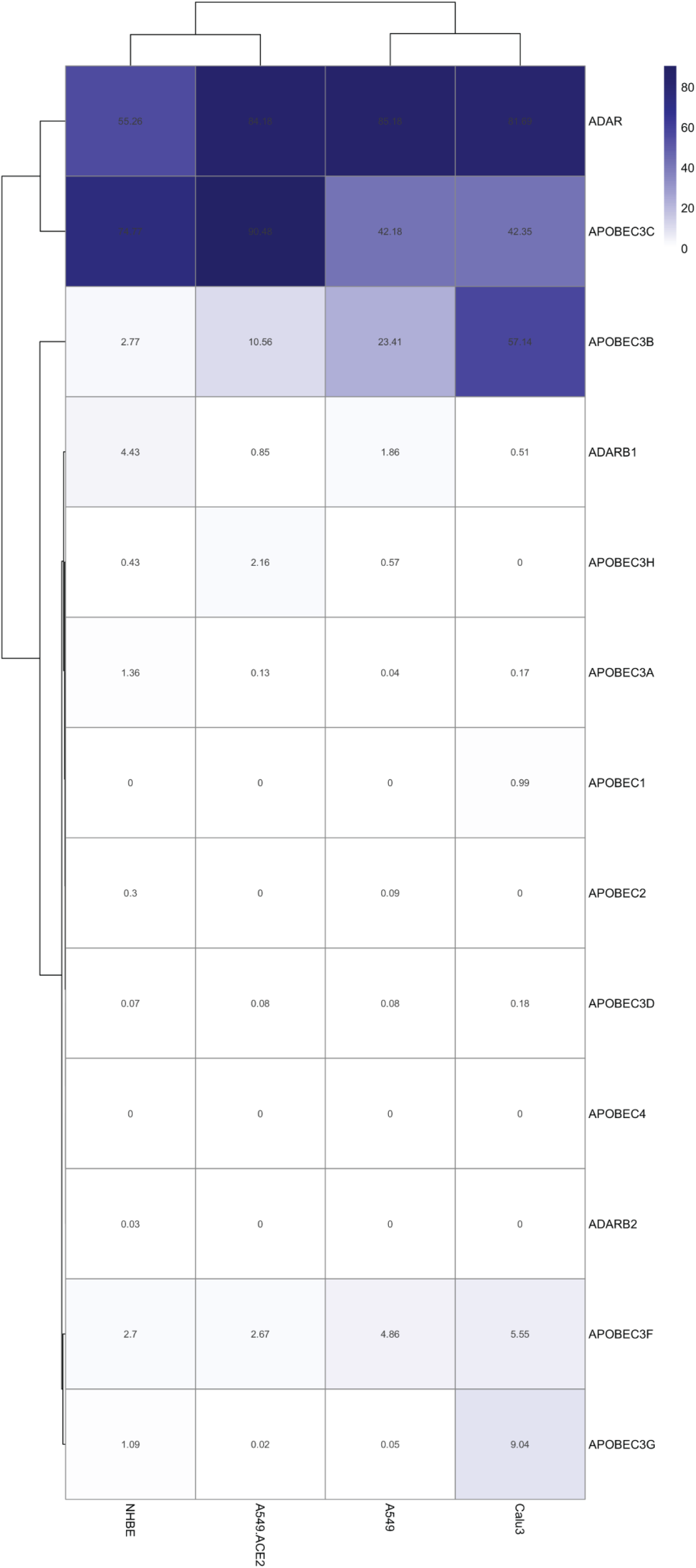
Heatmap representation for the expression of genes coding for APOBEC and ADAR family members. Counts are represented as Transcripts Per Million (TPM). The blue scale also correlates with TPM values.

**Fig. 4:**
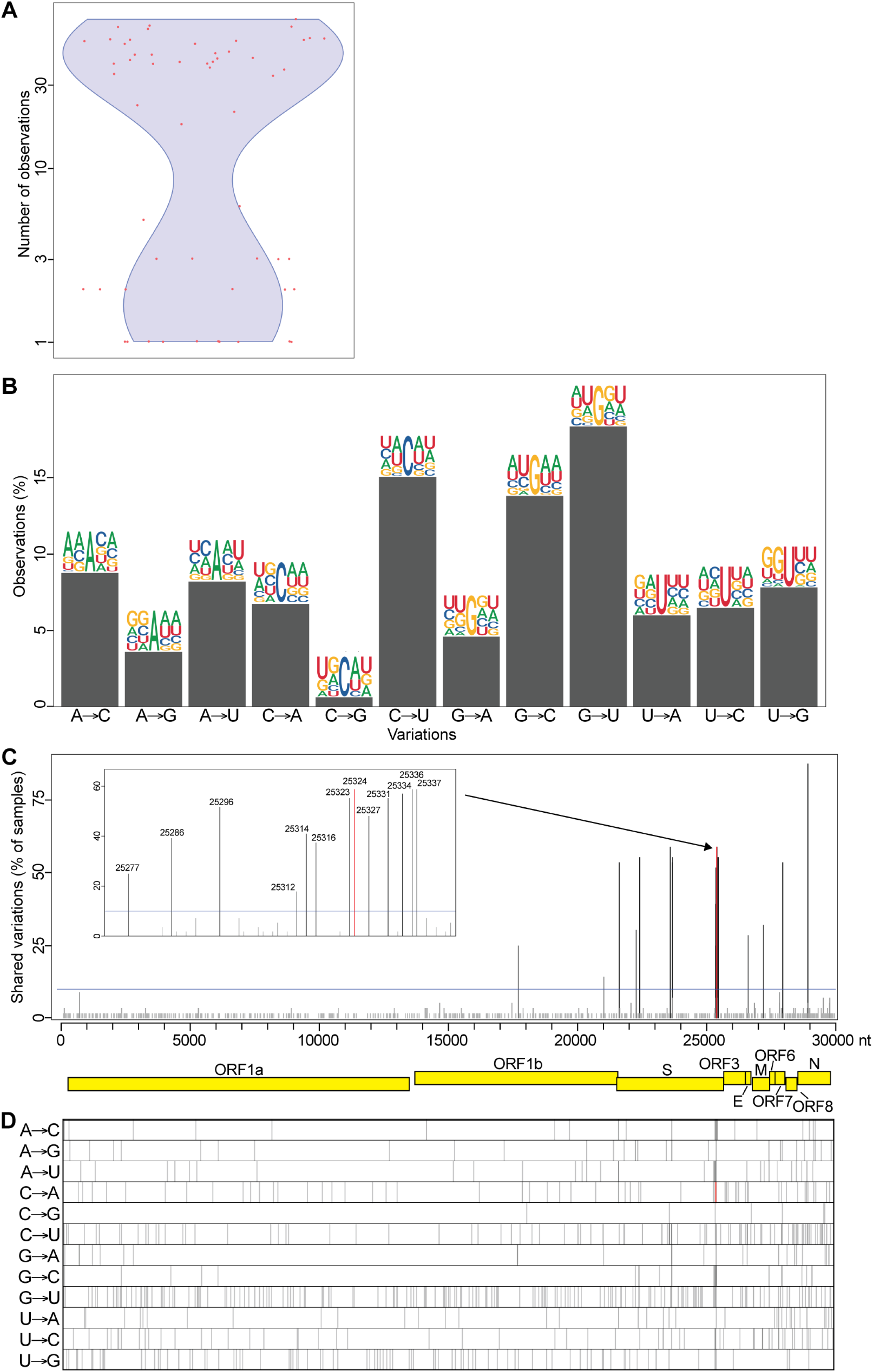
Intra-sample variability of the SARS-CoV-2 genome in infected cells. (**A**) Number of intra-variations observed for each sample analyzed. The red dots represent the 11,362 samples analyzed and the blue violon shows the distribution of the data. (**B**) Type of variation and sequence context for each intra-sample variable position. Bars represent the percentage of each type. Sequence context is represented by logos comprised of the consensus nucleotides (center) with two nucleotides upstream and two downstream from each intra-sample variable position. (**C**) Recurrent intra-genetic variations are represented as percentages of samples containing variation at each position. The SARS-CoV-2 genome and its genes are represented by yellow boxes below the graph. The blue line indicates 10% shared variations and was used to extract the intra-sample variation listed in Table 2. The inset represents a magnification of the cluster identified at the end of the S gene. (**D**) One-dimension representation of the data shown in panel C for each type of variation individually. The location of the C->A variation at position 25,324 is indicated by a red line in (C) and (D).

We then examined the intra-genetic variable positions that are recurrent amongst the cell lines analyzed. We identified 29 positions within the viral populations showing intra-genetic variation enrichment in at least 10% of the cell cultures and most of them are located on structural genes, which are encoded at the last 3’ terminal third of the viral genome (Fig. 4C and 4D). Similar to our observation from the samples from infected individuals, a cluster of recurrent variations is located at the 3’end of the S gene, including the C->A transversion at position 25,324 shared in 58.9% of the cell lines analyzed (Fig. 4C and 4D, red line; Table 2). Overall, our results indicate consistent results between intra-genetic variations observed in infected cell lines and in samples from infected individuals, including the presence the viral subspecies resulting in a S protein truncated of its last 20 amino acids (SΔ20).

**Table 2:**
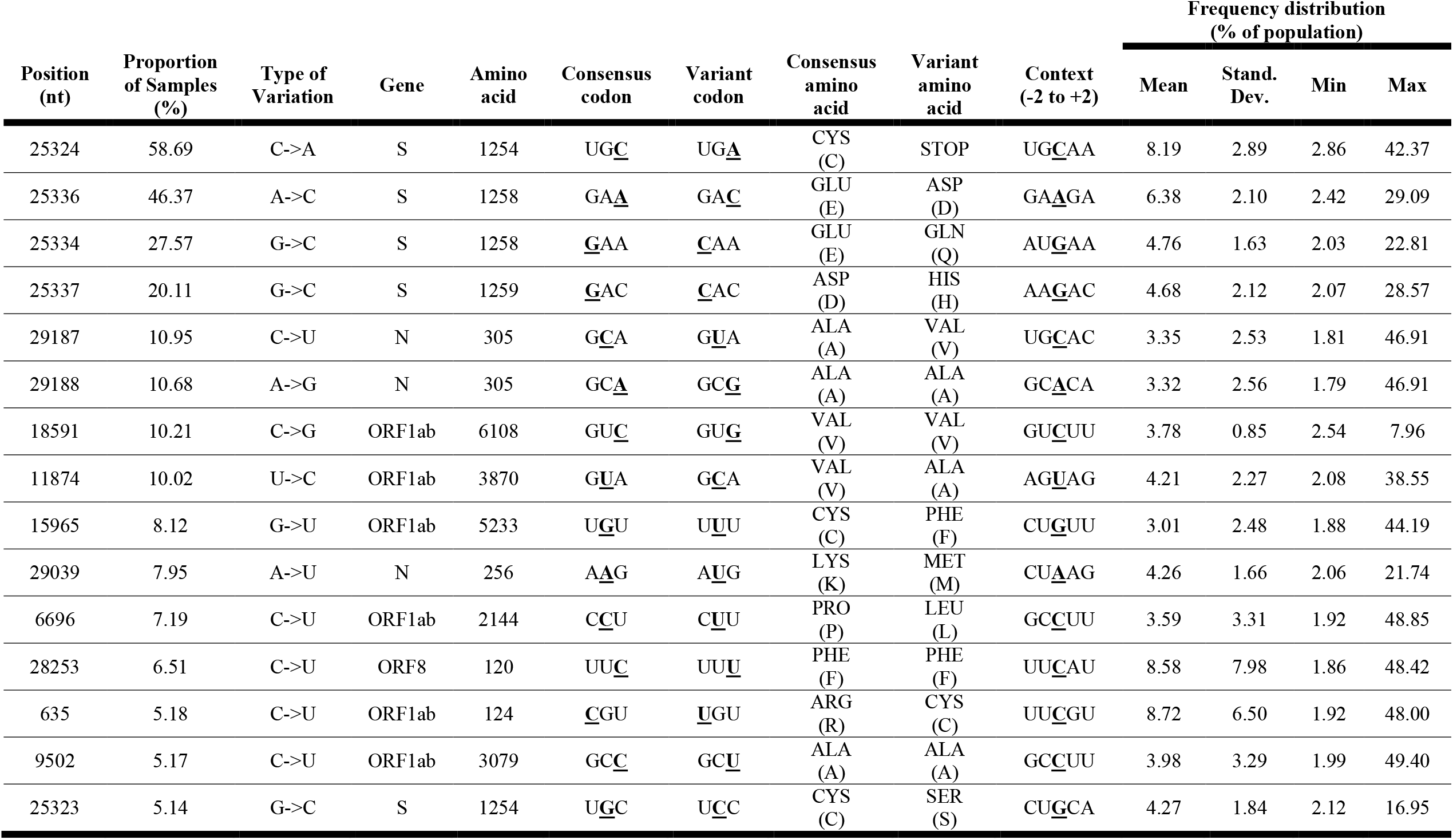
Recurrent SARS-CoV-2 genome intra-variations shared by at least 10% infected cell cultures. Frequency distributions were calculated using data similar to Fig. 2C. The variations are sorted by their recurrence, with the most shared variation at the top of the table.

### Increased fusogenic properties of SARS-CoV-2 SΔ20

SARS-CoV-2 viral entry into cells is triggered by the interaction between the S glycoprotein and its cellular receptor ACE2. While the complete mechanism of viral entry is not fully understood, it is known that S undergoes different processing steps by cellular surface and endosomal proteases. For several coronaviruses, the S protein not only mediates virion fusion but also syncytia formation (8, 13, 14). The presence of dysmorphic pneumocytes forming syncytial elements is well-described feature of the COVID-19 disease severity (37). One particularity of SARS-CoV-2 compared to SARS-CoV, is the presence of an additional furin-like cleavage site at the S1/S2 interface. As a consequence, SARS-CoV-2 cells have higher propensity to express activated S at the surface which can fuse with other cells expressing the receptor ACE2 and form syncytia (37). The normal route of S trafficking involves an accumulation at the ERGIC which is known to involve, at least in part, the interaction of the cytoplasmic portion of S with the M protein encoded by SARS-CoV-2. This interaction allows complex formation leading to virion formation at the ERGIC interface. The discovery of the SΔ20 variant missing a portion of the C-terminus directed us to investigate the effect on cell fusion using a syncytia assay in the presence of the M protein. HEK293T cells stably expressing the human ACE2 were co-transfected with plasmids encoding GFP, the M protein and the WT or Δ20 S protein. Consistent with previous findings (8), we observed syncytia formation in the presence of the S WT and SΔ20, indicating induction of cell-to-cell fusion (Fig. 5A). We also observed larger syncytia formation with SΔ20 compared to S WT, which indicates increased fusogenic activity of this truncated variant. As expected, the co-expression of the M protein and WT S completely abolishes syncytia formation which is a consequence of S being retained to the ERGIC. Strikingly, M protein failed to inhibit syncytia formation in the presence of the SΔ20 (Fig. 5A). To evaluate the effect of the Δ20 truncation on spike protein processing, we co-expressed the M protein with WT or Δ20 S protein in HEK293T in the absence of ACE2 to avoid cell fusion. Cells were lysed 24 hours post-transfection and spike processing was assessed by probing for SARS-CoV2 S1 and S2 subunits by immunoblotting. As seen in figure 5B, the SΔ20 undergoes increase processing as observed by the presence of more S1 and S2 compared to S WT (Fig. 5B; lane 2 vs lane 4). Finally, the co-expression of the M protein reduces the processing of the S WT protein while not affecting SΔ20 processing, as observed by a reduction of S1 fragment only for S WT (Fig. 5B; lane 3 vs lane 5). Taken together, our results indicate that SΔ20 displays increased processing and syncytia formation as compared to the wild-type S protein and the truncation removes an important regulatory domain involving the M protein.

**Fig. 5:**
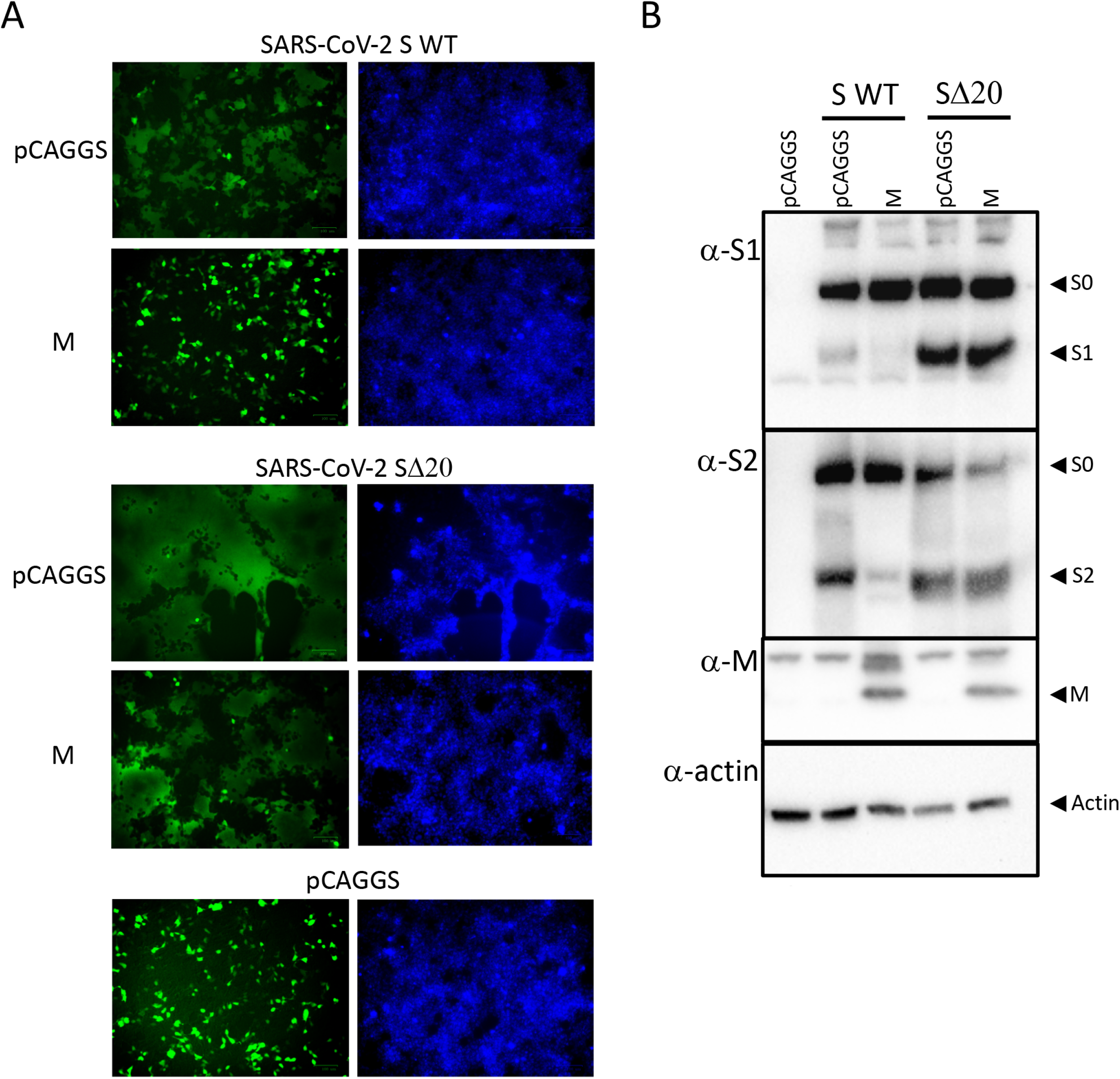
Increase processing and cytopathic syncytia formation by the SARS-CoV-2 spike Δ20. (**A**) Fluorescence microscopy of HEK-293T-hACE2 cells expressing GFP (green) with empty vector (pCAGGS), or plasmid expressing SARS-CoV-2 S or SARS-CoV-2 SΔ20 in the presence or absence of M protein. Counterstaining using Hoechst dye (blue) which labels nuclear DNA, is shown on the right panel. (B) Processing of spike was detected using an anti-S1 and anti-S2 immunoblotting of HEK293T cells lysates previously transfected with empty vector (pCAGGS), or vector expressing SARS-CoV-2 S or SARS-CoV-2 SΔ20 in the presence or absence of M protein.

## DISCUSSION

Previous analyses of SARS-CoV-2 nucleotide variations indicated a high prevalence of C->U transitions, suggesting that the viral genome was actively evolving and host editing enzymes, such as APOBECs and ADARs, might be involved in this process (24, 25). Although instructive on the role of host involvement in SARS-CoV-2 genome evolution, these studies were performed on consensus sequences (*i*.*e*., one per sample) and explore only part of the genetic landscape of this RNA virus. Here, we used a large number of high-throughput sequencing datasets to profile the intra-sample sequence diversity of SARS-CoV-2 variants, both in infected individuals and infected cell lines. We observed extensive genetic variability of the viral genome, including a high number of transversions, and identified several positions with recurrent intra-variability in the samples analyzed. Notably, most of the samples possessed a C->A missense mutation, producing an S protein that lacks the last 20 amino acids (SΔ20) and results in increased cell-to-cell fusion and syncytia formation.

Most intra-sample variations are distributed homogeneously across the viral genome, are not conserved or recurrent amongst samples, and a large number of them are C->U or G->U mutations. Previous analyses of SARS-CoV-2 sequence variations proposed that host editing enzymes might be involved in coronavirus transition editing, based on results showing that C->U transitions occur within a sequence context reminiscent of APOBEC1-mediated deamination (*i*.*e*., [AU]C[AU]) (22–25). Here, we investigated nucleotide compositions at each variation site and observed a high number of As and Us around all variation types and sites. However, since the SARS-CoV-2 genome is 62% A/U-rich and similar percentages of As and Us were observed around all variations, we concluded that no motifs are enriched around these variations in the viral subspecies analyzed. Consequently, our results cannot support that host editing enzymes are a major source of these intra-sample variations.

Although it is possible that host RNA-editing enzyme are responsible for the occurrence of some variations, C->U transitions and G->U transversions are also generally associated with nucleotide deamination and oxidation, respectively (38–45). It is common practice to thermally inactivate SARS-CoV-2 samples before performing RNA extractions, RT-PCR, and sequencing (46). However, heating samples can result in free radical formation, such as 8-hydroxy-20-deoxyguanine (8-Oxo-dG), that could cause high levels of C->A and G ->U mutations and promote the hydrolytic deamination of C->U (38–41, 43, 45, 47, 48). It was previously reported that these types of mutations occur at low frequency, that they are mostly detected when sequencing is performed on only one DNA strand, and that they are highly variable across independent experiments (40, 42). Consequently, most transversions observed in our analysis are likely due to heat-induced damage, RNA extraction, storage, shearing, and/or RT-PCR amplification errors. However, we identified several positions with intra-sample variability recurrent in several independent samples, both from infected individuals and infected cells. They were detected at moderate to high frequencies, ranging from 2.5% to 39.3% per sample (Table 1 and 2), and most were derived from pair-end sequencing (90.7%) in which the two strands of a DNA duplex were considered. Thus, it is likely that these variations are genuine and represent hot spots for SARS-CoV-2 genome intra-sample variability.

Amongst the variable positions identified in infected cells, most of them are located at the last 3’ terminal third of the viral genome. These cells were infected with a large number of viruses (*i*.*e*., a high multiplicity of infection; MOI) for 24h (30). The presence of several variations at positions in the region coding for the main structural proteins likely reflects that this is a region with increased transcriptional activity due to the requirement of producing their encoded mRNAs from sub-genomic negative-sense RNAs (9).

Interestingly, a cluster of variations located at the 3’end of the S gene was observed for the two datasets analyzed. They correspond to four transversions located at the 3’end of the S gene and are shared by a large proportion of the samples. Three of these correspond to missense mutations changing the charged side chains of two amino acids (E1258D, E1258Q and D1259H). Notably, most of the samples possess a variability at position 25,324, producing a nonsense mutation at amino acid 1,254 of the S protein. The resulting protein lacks the last 20 amino acids (SΔ20) and thus does not include the ERRS motif at its carboxy-terminus. For SARS-CoV-1, the ERRS domain accumulates the S protein to the ERGIC and facilitates its incorporation into virions (12). While the mechanism is not completely understood, mutation of the ERRS motif on S resulted in a failure to interact with the M protein at the ERGIC and rather resulted in trafficking of S to the cell surface. Deletion of this motif might cause the S protein of SARS-CoV-2 to accumulate to the plasma membrane and increase the formation of large multinucleated cells known as syncytia. Consistent with these observations, our results indicate larger syncytia formation with SΔ20 compared to the complete S protein. Moreover, we observed that the M protein failed to prevent SΔ20-induced syncytia formation, as observed with the WT S protein, which correlates with the role of the M protein in interacting with Spike and retaining it in ERGIC. Similar mutants (SΔ18, SΔ19 and SΔ21) were recently reported to increase both infectivity and replication of vesicular stomatitis virus (VSV) and human immunodeficiency virus (HIV) pseudotyped with SARS-CoV-2 S protein in cultured cells (49–52). Because these viruses bud from the plasma membrane (53, 54), an increased localization at this site would explain the selection of these deletion mutants in pseudotyped virions. However, such variants would unlikely be transmitted horizontally in naturally occurring CoV where the budding site is the ERGIC compartment (10).

Our findings indicate the presence of consistent intra-sample genetic variants of SARS-CoV-2, including a recurrent sub-population of SΔ20 variants with elevated fusogenic properties. It is tempting to suggest a link between SARS-CoV-2 pathogenesis and the presence of SΔ20, since severe cases of the disease were recently linked to considerable lung damage and the occurrence of syncytia (37, 55). Clearly, more investigation is required to better define the extent of SARS-CoV-2 variability in infected hosts and to assess the role of these subspecies in the life cycle of this virus. More importantly, further studies on the presence of SΔ20 and its link with viral pathogenicity could lead to better diagnostic strategies and design treatments for COVID-19.

## ACKNOWLEDGEMENTS

K.F. is supported by an Ontario Graduate Scholarship. C.M.S. is supported by a graduate scholarship from the Natural Sciences and Engineering Research Council of Canada. M.-A.L. holds a Canada Research Chair in Molecular Virology and Intrinsic Immunity. M.C. is a Canada Research Chair in Molecular Virology and Antiviral Therapeutics. This work was supported by a COVID-19 Rapid Research grant from the Canadian Institutes for Health Research (CIHR; OV1 170355) to M.-A.L and M.P., and a COVID-19 Rapid Research Grant (OV3 170632) to M.C. and P.M.G.

